# In situ protein micro-crystal fabrication by cryo-FIB for electron diffraction

**DOI:** 10.1101/377192

**Authors:** Xinmei Li, Shuangbo Zhang, Fei Sun

## Abstract

MicroED (micro electron diffraction) is an emerging technique to use cryo-electron microscope to study the crystal structure of macromolecule from its micro/nano-crystals, which are not suitable for conventional X-ray crystallography. However, this technique has been prevented for its wide application by the limited availability of producing good micro-/nano-crystals and the inappropriate transfer of crystals. Here, we developed a complete workflow to prepare suitable crystals efficiently for MicroED experiment. This workflow includes *in situ* on-grid crystallization, single-side blotting, cryo-focus ion beam (cryo-FIB) fabrication, and cryo-electron diffraction of crystal cryo-lamella. This workflow enables us to apply MicroED to study many small macromolecular crystals with the size of 2 ~ 10 μm, which is too large for MicroED but quite small for conventional X-ray crystallography. We have applied this method to solve 2.5Å crystal structure of lysozyme from its micro-crystal within the size of 10×10×10 μm^3^. Our work will greatly expand the availability space of crystals suitable for MicroED and fill up the gap between MicroED and X-ray crystallography.

## INTRODUCTION

In 1960, the crystal structures of myoglobin and hemoglobin were solved (Kendrew *et al*, 1960; Perutz *et al*, 1960a; Perutz *et al*, 1960b) by X-ray crystallography, opening the era of structural biology. Till now, there have been hundreds of thousands of bio-macromolecular structures that were determined by X-ray crystallography. The size of bio-macromolecular crystal should be large enough to gain efficient signal-noise ratio (SNR) of diffractions. For the home source of X-ray generated from rotating anode, the size of crystal needs to be larger than 200 μm normally (Mizohata *et al*, 2018). While the emerge of synchrotron radiation allows a brilliant and coherent source of X-ray, which can increase SNR of diffractions especially at the high resolution region, thus even a smaller crystal (ca. 50 ~ 200 μm) still generate significant diffractions for structure determination (Mizohata *et al*, 2018). The emerge of the micro-focus beamline based on the third-generation synchrotron source has yield micro-crystallography, which made high-resolution data collection from very small crystals (ca. 10 ~ 50 μm) possible (Smith *et al*, 2012). The widespread use of synchrotron radiation has accelerated the development of X-ray crystallography and structural biology.

However, many bio-macromolecules could not be crystallized into large crystals, e.g. membrane proteins and bio-macromolecular complexes. When the size of crystal is further smaller than 10 μm, the current source of X-ray from synchrotron radiation could not yield high SNR diffractions. More importantly, the severe radiation damage from high flux X-ray exposure makes it impossible to collect a complete diffraction dataset from a single crystal, even under the cryogenic condition (Johansson *et al*, 2017). The emerge of X-ray free electron laser (XFEL) and the development of serial femtosecond X-ray crystallography (SFX) provide an alternative solution (Chapman *et al*, 2011). The extremely short and highly intensive X-ray pulse makes it possible to collect high SNR and ‘radiation damage free’ diffraction data from a single microcrystal (ca. 1 ~ 10 μm) (Mizohata *et al*, 2018). Each microcrystal generates one frame of diffraction image before obliterated (Neutze *et al*, 2000). Tens of thousands of microcrystals are needed to generate a complete dataset. In recent years, SFX has been successfully applied to solve many important and difficult crystal structures, including the human Angiotensin II type 1 (Zhang *et al*, 2015) and photon-synthesis complex II (Suga *et al*, 2017). Furthermore, the extremely short pulse (10 ~ 100 fs) of XFEL also enables time-resolved SFX and people can investigate the transient structural changes of photon-synthesis complex II upon light stimulation (Suga *et al*, 2017). However, culturing tens of thousands of microcrystals and the limited accessibility of XFEL facility have restricted the wide application of SFX technology.

Besides diffracting X-ray, bio-macromolecular crystals could also diffract electron, which is called electron crystallography when 2D crystals are investigated, or called micro-electron diffraction (MicroED) for 3D crystals. Investigation on biological specimens by electron crystallography arguably began when Parsons and Martius used electron diffraction to investigate the structure of muscle fibers in 1964 (Parsons and Martius, 1964). In 2005, the 2D electron crystallography has been successfully utilized to solve the high resolution structure of water channel AQP0 in a closed conformation at 1.9 Å (Gonen *et al*, 2005). Considering there are only 17 plane groups allowed for 2D protein crystals while there are 65 symmetries allowed for 3D protein crystals (Nannenga Brent *et al*, 2013), it is of less successful rate to grow a 2D crystal rather than a 3D crystal (Martynowycz and Gonen, 2018). Furthermore, the extreme difficulty of culturing high-quality 2D crystals of bio-macromolecules has limited the wide application of 2D electron crystallography.

In 2013, Shi *et al*. first utilized MicroED to solve the crystal structure of lysozyme at 2.9 Å resolution (Shi *et al*, 2013) and later improved the data quality by changing still diffraction mode to continuous rotation diffraction mode (Nannenga *et al*, 2014b). Compared with X-ray, electron has much larger scattering cross-section when interacting with atoms (Henderson, 2009). Thus, the size of microcrystal is large enough to generate high SNR diffractions by using MicroED. The size of crystal used in MicroED experiment is within 500 nm (Nannenga and Gonen, 2014), much smaller than the one in traditional X-ray crystallography experiments.

In recent years, MicroED has been further developed and applied to solve the crystal structures of a-synuclein (Rodriguez *et al*, 2015), prions (Sawaya *et al*, 2016) and the human fused in sarcoma low-complexity domain (FUS LC) (Luo *et al*, 2018) in atomic resolution. The crystals of these successful examples were always thin although they were long and wide. For example, the crystals of the a-synuclein are needle-like with the thickness of 20 ~ 50 nm (Rodriguez *et al*, 2015). The thickness of the crystal directly determines the quality of diffraction data (Nannenga and Gonen, 2014) due to the mean free path of electron. For 300 kV electron, its mean free path for the vitrified bio specimen is ~ 350 nm, while for 200 kV electron, it is ~ 300 nm (Yan *et al*, 2015). When the thickness of the crystal is over beyond the mean free path of electron, multi-scattering events will become significant and then the diffraction pattern will become difficult to explain. As a result, for a success of MicroED experiment, nanocrystals not microcrystals are actually needed.

The emergence of MicroED has provided an alternative solution to X-ray crystallography and it is possible to solve the crystal structure of bio-macromolecule using a single nanocrystal by MicroED. However, there are still several bottlenecks left, limiting the wide application of MicroED. Firstly, it is difficult to grow and screen nanocrystals. Nanocrystals are invisible under light microscope and their growth process is difficult to monitor. Transmission electron microscope is the only way to screen the presence of nanocrystals, which is in a low throughput. The previous trials of breaking big crystals into tiny bricks were not successful (personal communication). Secondly, the current sample freezing procedure (blotting and plunge-freezing) for single particle analysis is not optimized for MicroED sample preparation. The viscous crystallization liquid could not be easily blotted and the double-sided blotting step could damage delicate crystals easily (Shian *et al*, 2017). In addition, more importantly, there are many bio-macromolecules that can be crystallized into microcrystal with the size of 2 ~ 10 μm. These crystals could not be analyzed by traditional X-ray crystallography and even by SFX easily. For MicroED, the size of these crystals is also beyond the feasible range.

The recent emergence of cryo focused ion beam (cryo-FIB) technique (Marko *et al*, 2006) provides a solution to prepare suitable size of crystal for MicroED experiment. This technique was first applied to prepare a cryo-lamella of bacterial cells (Marko *et al*, 2007) and later to eukaryotic microbial cells (Rigort *et al*, 2012) and mammalian cells (Strunk *et al*, 2012); (Wang *et al*, 2012). We also developed this technique with the name of D-cryo-FIB (Zhang *et al*, 2016), which has been used to prepare cryo-lamella for subsequent cryo-electron tomography experiment (Li *et al*, 2015).

Here, based on our D-cryo-FIB technique, we report a workflow of *in situ* protein crystallization and fabrication for subsequent successful MicroED experiment. We grew protein crystals on grids directly to reduce the possibility of crystal missing during sample transfer. The grids were blotted from back side to alleviate the crystal damage due to blot force. Then the crystals frozen on the grid were selected and milled into a thin lamella by cryo-FIB. The cryo-lamella was then used to collect electron diffraction dataset for structure determination. We show here that we successfully solved the crystal structure of lysozyme at 2.5 Å resolution by using seven micro lysozyme crystals within the size of 10×10×10 μm^3^.

## MATERIALS AND METHODS

### *In situ* crystallization and cryo-vitrification

Lysozyme was purchased by Sangon Biotech Company. A 200 mg/ml solution of lysozyme was prepared in 50 mM sodium acetate pH 4.5. A grow-discharge treated nonmagnetic nickel grid with holy carbon film was placed facing up on a micro-bridge, and the protein was mixed 1:1 with the precipitant solution (0.35 M sodium chloride; 15% PEG5000MME; 50 mM sodium acetate pH 4.5) on the grid. Then the crystals were grown by the sitting drop vapor diffusion method (**Fig. 1**).

**Figure 1.**
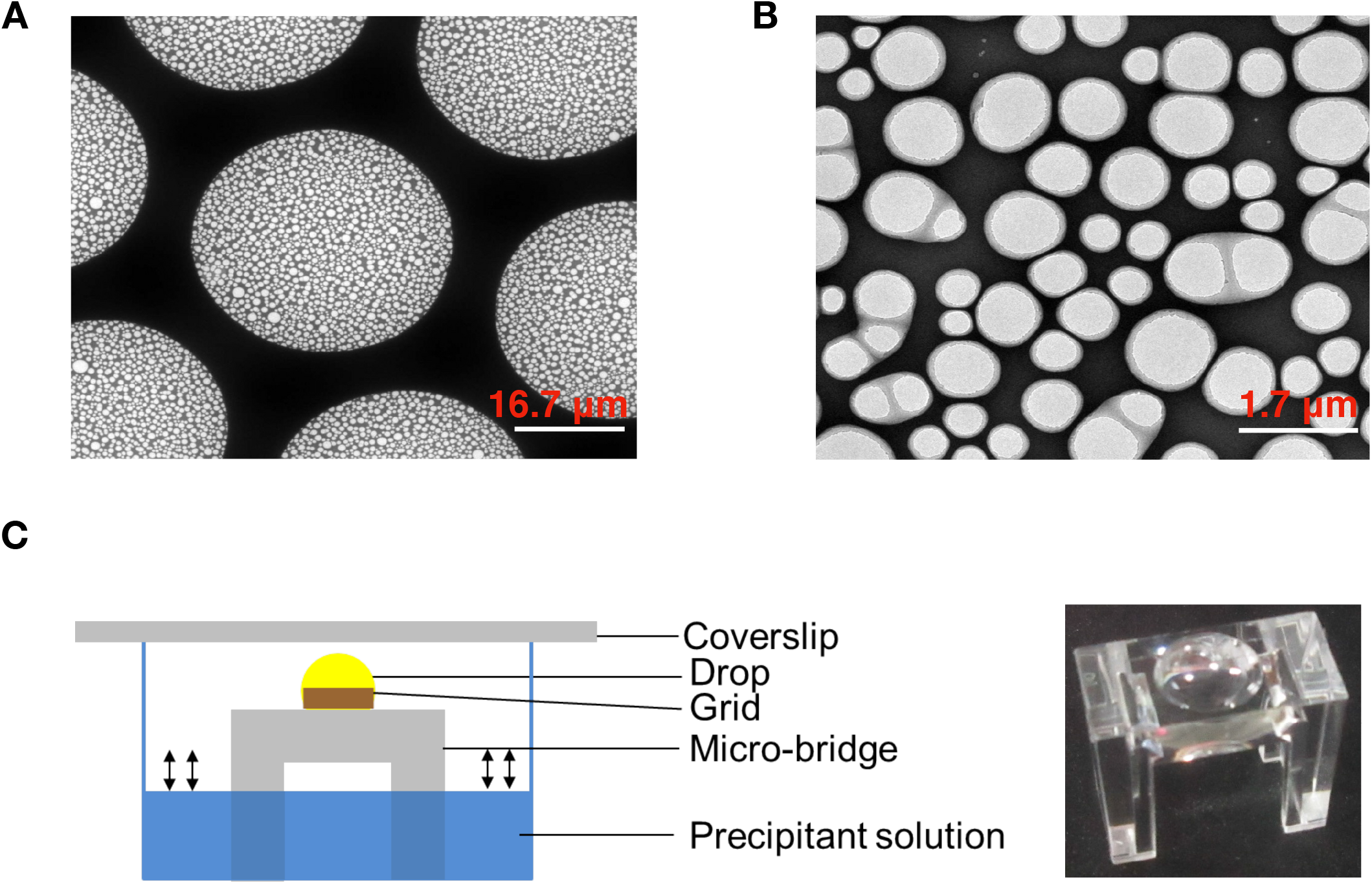
*In situ* on-grid protein crystallization setup. (A) A magnified SEM image of the nonmagnetic nickel grid coated with holy carbon film. Scale bar, 16.7 μm. (B) A further magnified SEM image in (A) showing the structural details of the holy carbon film. Scale bar, 1.7 μm. (C) The schematic diagram of the sitting drop vapor diffusion setup for *in situ* on-grid protein crystallization. The real photo of the micro-bridge is shown in right. Scale bar, 0.35 mm.

After the crystals formed, the grid was washed four times by washing buffer (0.35 M sodium chloride; 5% PEG200; 50 mM sodium acetate pH 4.5) before cryo-vitrification. Then the grid was blotted 6 s from the backside where no crystals grew on, and plunged into liquid ethane using the Leica EMGP. The frozen grids were transferred and stored in liquid nitrogen for the subsequent experiments.

### Crystals fabrication by cryo-FIB

The frozed grid was loaded onto a home-made cryo-shuttle (Zhang *et al*, 2016) and then transferred into the chamber of a dual beam scanning electron microscope (FEI Helios NanoLab 600i) that is equipped with a Quorum PT3000 cryo-stage. The grid has been pre-tilted 45° on the shuttle. Then the grid was imaged and examined using electron beam and the signal from secondary electron with the following experimental parameters, an accelerating voltage of 2 kV, a beam current of 0.34 nA, and a dwell time of 3 μs. The region of interest (ROI) with crystals was identified and marked. Then focused ion gallium beam was utilized to perform fabrication of crystals. Before cryo-FIB milling, a thin layer of Pt was coated using GIS system to reduce the radiation damage during milling. The inclination angle of the beam was kept ~ 10° against the grid plane. The accelerating voltage of ion beam was 30 kV. Considering the block shape of the crystal, a large beam current of 0.43 nA was used to skive crystals efficiently. When a thick slice of lamella was made, the beam current was reduced to 80 pA for the fine trimming and also the reduction of potential radiation damage. The final thickness of the crystal cryo-lamella was controlled ~ 300 nm. Several crystal cryo-lamellas can be made in one grid. After cryo-FIB fabrication, the grid was transferred and kept in liquid nitrogen for the subsequent MicroED experiment.

### Cryo-electron diffraction and data collection

Cryo-electron diffraction of cryo-FIB fabricated crystal was collected using cryo-electron microscope FEI Talos F200C equipped with a field-emission gun operated at 200 kV (λ= 0.0251 Å). And the diffraction patterns were recorded by the FEI Ceta camera with 4096×4096 pixels and the physical pixel size of 14 μm. We utilized SerialEM (Mastronarde, 2005) to control the microscope and collect the diffraction datasets.

The frozen grid was loaded into the microscope using a Gatan cryo-transfer holder (Model 626) that was precooled in liquid nitrogen. The straight side of the D-shaped grid (Zhang *et al*, 2016) was kept parallel to the rotation axis of the holder. ROIs that were trimmed by cryo-FIB were located in low magnification of SA2600X (view mode in SerialEM).

Then, at the exposure mode in SerialEM, the spot size and the illumination area were adjusted to yield a very low electron dose of 0.07 e^-^/Å^2^/s for the diffraction experiment. To measure the electron dose accurately, the microscope should be in image mode and high magnification so that the electron beam can spread over the entire screen. Eventually, in our experiment setup, the spot size was selected as 9 and the excitation level of C2 lens was kept at 45.3000%. After the electron dose was determined, the microscope was switched to diffraction mode. The nominal camera length was set to 1.35 m that was calibrated to 2.22 m by measuring the diffraction pattern of gold crystal. The excitation level of objective lens was set and kept at 85.5148%. The diffraction lens was adjusted to focus the central beam. Then, we switched back and forth between the image and diffraction modes to make sure that the excitation levels of objective and C2 lens did not change. Finally, we switched to diffraction mode and saved all the parameters to the exposure mode in SerialEM.

When all the parameters were setup, we moved the cryo-FIB fabricated crystal to the center of the screen in the view mode of SerialEM, and then chosen a selective aperture of 100 μm in diameter to just cover the area of the crystal. Then we switched to the exposure mode in SerialEM. A customized SerialEM script (see **Supplementary Information**) was written to collect diffraction data in an approximate continuous rotation mode, in which the crystal was initially rotated to −40° (or other degree) and then started to rotate to 40° (or other degree). For every 0.2° increment, a diffraction frame was recorded with the exposure time of 0.2 s and stored in TIFF format. For the rotation angle range from −40° to 40°, we collected 400 frames with the approximate total dose of ~6 e^-^/Å^2^/s. In total, we collected multiple diffraction datasets from different cryo-FIB fabricated crystals. For each crystal, the total electron dose was kept below 9 e^-^/Å^2^/s.

### Processing of electron diffraction datasets

We first exacted the dark background from every raw diffraction image and then summed every five frames to the final one with an “expected” oscillation angle width of 1°. The program developed in Tamir Gonen’s lab for converting TVIPS camera image to the SMV format (http://cryoem.janelia.org/pages/MicroED) was slightly modified based on our microscope and camera system, and then used to convert our datasets from TIFF format to SMV format.

Then the converted diffraction datasets were processed (index and integration) by iMOSFLM 7.2.1 (Battye *et al*, 2011). Different with processing X-ray crystallography datasets, a wide rotation angle range was used for a successful index because the Ewald sphere in electron diffraction is very flat. Secondly, due to the instability of microscope stage, the crystal orientation and the distance from the crystal to the detector would change during data collection, which should be carefully considered during data integration process. Furthermore, our camera FEI CETA was not well characterized for electron diffraction experiment, and the GAIN parameter defined in iMOSFLM is generally unknown. Thus, it is important and necessary to try different GAIN values (1.5 ~ 2.5) during data integration process.

Due to the instability of our microscope, the first several frames of each dataset were dropped. Seven datasets were merged, scaled and converted to structure factor amplitudes using AIMLESS (Evans and Murshudov, 2013) to increase the completeness of diffraction data. Then molecular replacement was performed using PHASER (McCoy *et al*, 2007) with the starting model (PDB code, 4AXT) of lysozyme structure. Finally, REFMAC5 (Murshudov *et al*, 2007) was used to perform structural refinement by taking electron scattering factors into consideration.

The statistics of data collection, processing and structural refinement were summarized in **Table 1**.

**Table 1.** Statistics of data collection, processing and structural refinement

|  |  |
| --- | --- |
| **Data collection** |  |
| Excitation voltage (kV) | 200 |
| Electron source | Field emission gun |
| Wavelength (Å) | 0.025 |
| Total electron dose per crystal ($e^-/\text{Å}^2$) | 5 ~ 9 |
| Number of patterns per crystal | 150 ~ 400 |
| No. crystals used | 7 |
| Nominal camera length (m) | 1.35 |
| Real (corrected) camera length (m) | 2.22 |
| Selective area aperture ($\mu\text{m}$) | 100 |
| Rotation step ($^\circ$) | 0.2 |
| **Data Processing** |  |
| Resolution (Å) | 18.4 ~ 2.5 |
| Space group | P4<sub>3</sub>2<sub>1</sub>2 |
| Unit cell dimensions |  |
| a = b (Å) | 77 |
| c (Å) | 37 |
| $\alpha = \beta = \gamma$ ($^\circ$) | 90 |
| No. Total reflections | 53,003 |
| No. Unique reflections | 4,235 |
| CC<sub>1/2</sub> (overall / outer shell) | 0.803 / 0.158 |
| $\langle I / \sigma \rangle$ | 5.5 |
| Completeness (%) (overall / outer shell) | 94.0 / 86.8 |
| Multiplicity (overall / outer shell) | 12.5 / 12.7 |
| R<sub>merge</sub> | 0.356 |
| **Structural refinement** |  |
| Resolution (Å) | 18.4 ~ 2.5 |
| Reflections in working set | 4,206 |
| Reflections in test set | 191 |
| R<sub>work</sub>/R<sub>free</sub> (%) | 35.9 / 40.0 |
| r.m.s.d. Bond Length | 0.005 |
| r.m.s.d. Bond Angle | 0.976 |

## RESULTS AND DISCUSSIONS

To set up a workflow from *in situ* protein crystallization, cryo-vitrification, and cryo-FIB fabrication to the subsequent cryo-electron diffraction data collection, we selected lysozyme as the testing sample since it is well characterized and was previously used for MicroED experiments (Shi *et al*, 2013).

Originally, we followed the previously published protocol (Shi *et al*, 2013) to grow lysozyme crystals by hanging drop vapor diffusion method, and tried to add the crystallization drop from the cover slip to the grid for 1 min adsorption before washing and vitrification. However, by examining in cryo-electron microscope, we found the low successful rate of the crystal absorption on the grid and the crystal could be easily destroyed by the pipette during the drop transfer process. Thus, to overcome this difficulty, we sought to try growing crystals directly on the grid to reduce the loss of the crystal and minimize its potential damage during sample transfer. A holy carbon coated metal grid (**Figs. 1A and 1B**) was used in the present study. The sitting crystallization drop was directly added to the surface of a grow-discharge treated grid that is supported by a clean plastic micro-bridge (**Fig. 1C**). This experimental setup allows the crystals to grow on the carbon surface of the grid.

Different metal grids were screened to select the most suitable one for *in situ* crystallization. For the copper grids, we found the copper is chemically reactive in crystallization buffer and the dissolved copper ion could denature the protein and affect the quality of the crystal severely (see the green colored crystal in **Fig. 2A**), which was subsequently proved by X-ray diffraction experiments (data not shown here). Then the titanium (**Fig. 2B**), molybdenum (**Fig. 2C**) and nonmagnetic nickel grids (**Fig. 2D**) were tested. All these three grids could allow crystal grow on the carbon surface. Since these grids are stainless with good chemical inertia, there were no chemical effects observed to decrease the crystal quality. Considering the cost and availability, we selected the D-shaped nonmagnetic nickel grid (**Fig. 2D**) for the subsequent experiment.

**Figure 2.**
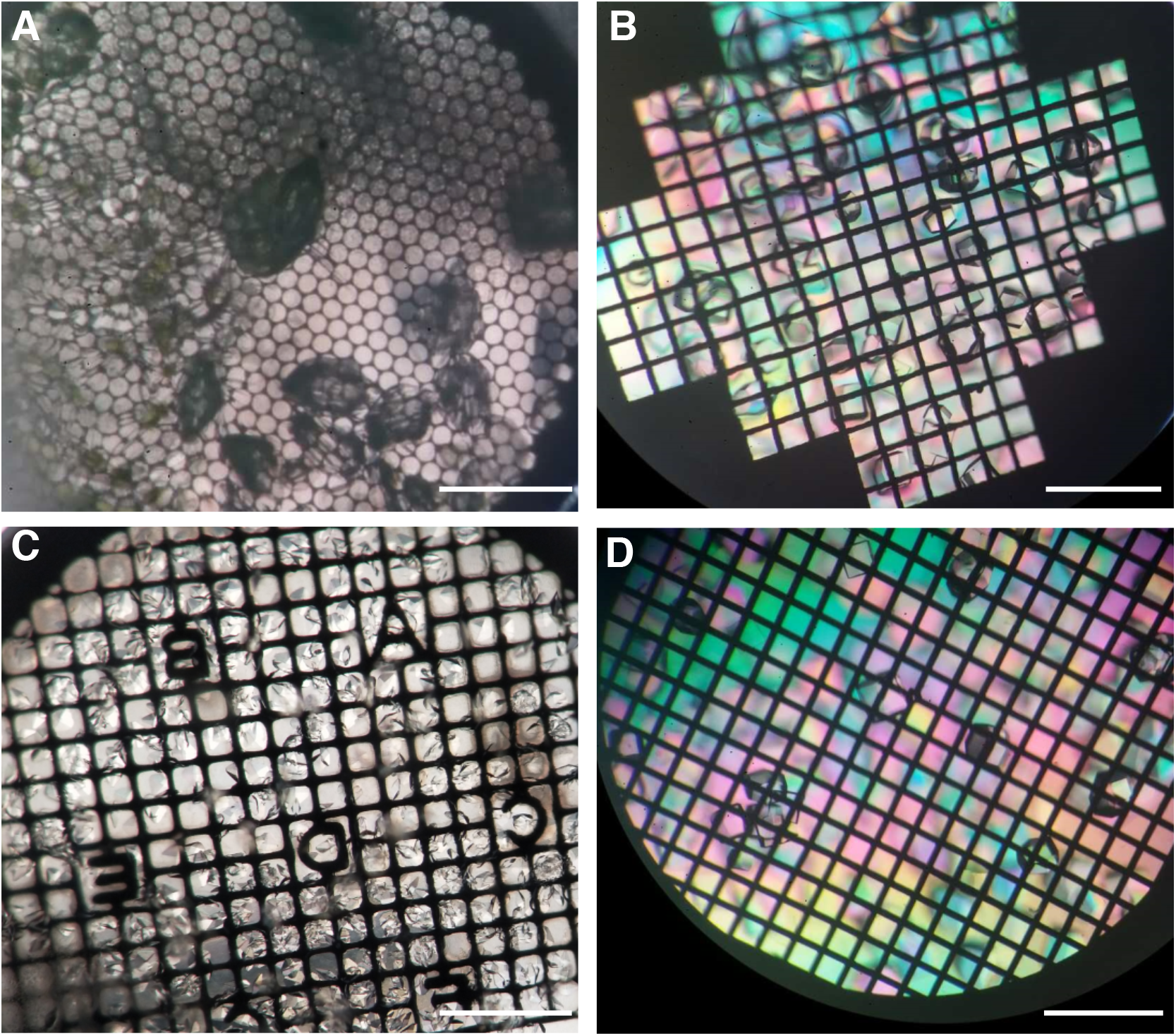
Photographs of the lysozyme crystals growing on different grids. (A) Copper grid. (B) Titanium grid. (C) Molybdenum grid. (D) Nonmagnetic nickel grid. Scale bar, 500 μm.

The crystallization drop contains 15% PEG5000MME that has a high viscosity and is difficult to blot to get a thin layer during cryo-vitrification process. To overcome this difficulty, we used the washing buffer that contains 5% PEG200 instead of 15% PEG5000MME to wash the grid before blotting, which helped to get a thin ice layer on the grid after freezing. To optimize the washing buffer that does not affect the quality of the crystals, we used to pick out some crystals and transfer them to the washing buffer and observed under light microscope. There should be no obvious dissolving phenomenon appeared within one hour. After washing, the grid was blotted for 6 seconds from backside using Leica EMGP and then fast frozen in liquid ethane. The backside blotting is also important to prevent potential crystal loss and damage.

The vitrified D-shaped grid was transferred into the chamber of FIB/SEM by keeping the direction of the straight edge perpendicular to FIB (Zhang *et al*, 2016). The crystals were first identified and visualized by SEM (**Figs. 3A and 3B**). To choose the area of interest for FIB milling, there are a few criteria to be considered. Firstly, the position of the crystal should be close to the middle of the square. Otherwise, the metal grid bar would block FIB, causing the trimming process not thorough. In addition, the grid bar could also potentially block the electron beam during diffraction data collection when the grid is tilted. Secondly, it is important to select a separated single crystal to avoid potential twin diffraction images during data collection. In addition, a proper size (5 ~ 20 μm) of the crystal was selected. Smaller size would decrease the successful rate of the cryo-lamella production and also yield a very small area for MicroED. Larger size would increase significantly the time of FIB trimming and thus lower the throughput. After the cryo-lamella of crystal was formed, its thickness was measured from FIB image and could be judged from the low magnification TEM image (**Fig. 3C**). For a good crystal lamella, its electron diffraction could reach to 2.0 Å resolution (**Fig. 3D**) with our current experiment hardware.

**Figure 3.**
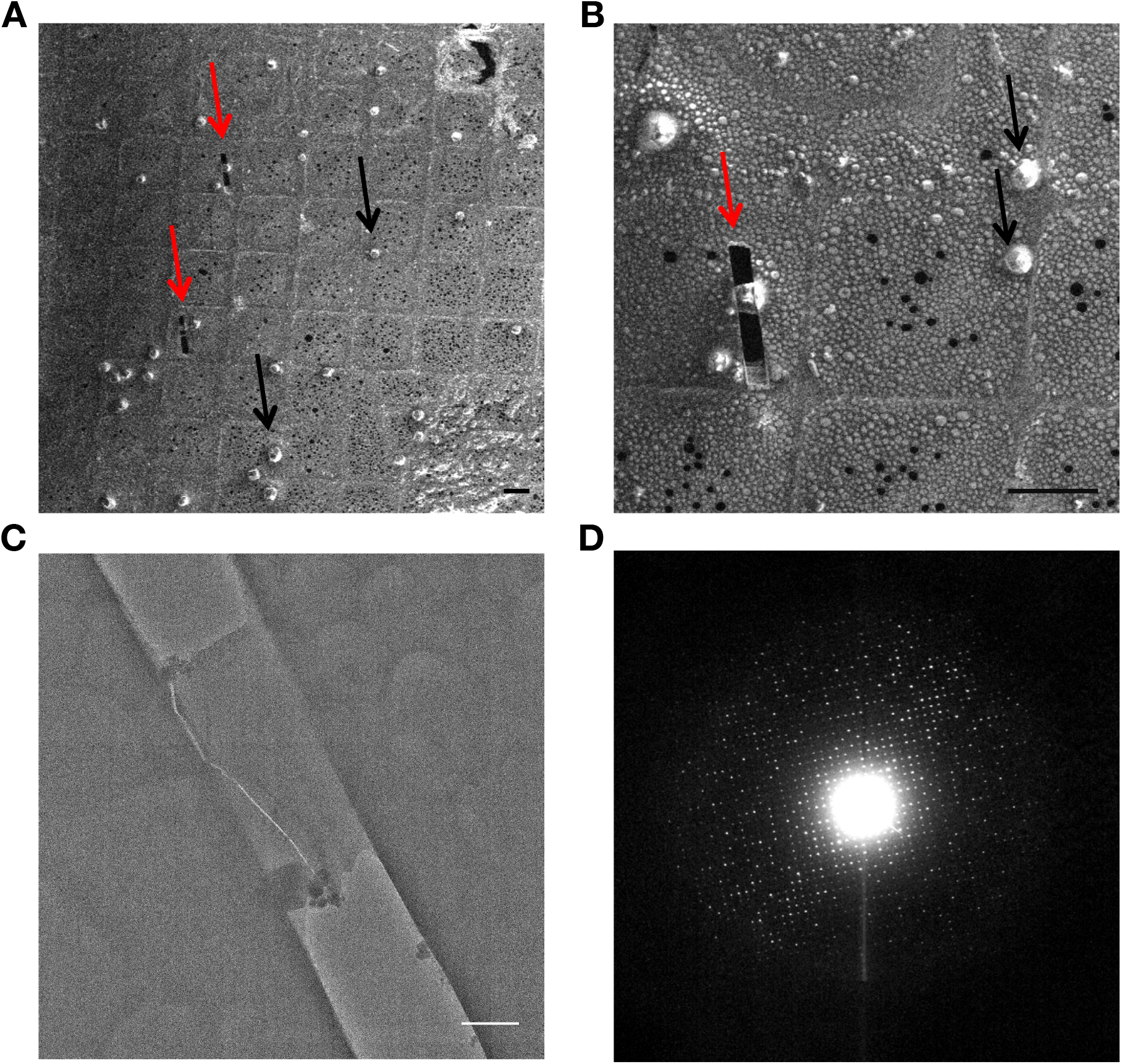
FIB fabrication of frozen lysozyme crystals on grid. (A) and (B) SEM images (SE detector) of frozen lysozyme crystals on grid with low magnification (A) and high magnification (B). The areas with strong contrast indicate the positions of the crystals. The black arrows indicate the crystals close to the grid bars and the red arrows indicate the crystals trimmed by FIB. Scale bars, 50 μm. (C) TEM micrograph of the FIB fabricated crystal lamella. Scale bar, 500nm. (D) Cryo-electron diffraction pattern of the FIB fabricated crystal lamella in (C).

During electron diffraction data collection, crystals could be either tilted discretely or rotated continuously in the electron beam, which yield still diffraction image or continuous diffraction image. We wrote a simple script of SerialEM (see **Supplementary Information**) to collect tilt series of still diffraction images automatically. We were aware of that collecting continuous diffraction images could increase the accuracy of reciprocal spot intensity measurement. However, for our current camera setup, it was difficult to synchronize the stage rotation with the data recording of camera due to the imperfect mechanics of the stage and the significant lag of camera recording system. Thus, we made an appropriate approach to collect continuous diffraction images. In our approach, we utilized our script to collect tilt series of still diffraction frames with a very small angle interval. In the present work, the angle interval of 0.2° was used. The exposure time for each still diffraction frame was properly determined to balance its SNR and the total electron dose, and was set 0.2 s/frame in the present work. Then, numbers of frames were simply integrated to form a final image that approximates the continuous diffraction one. Here, every five continuous frames were merged into one image with a rotation angle width of 1°, which was ready for the subsequent processing in iMOSFLM (**Fig. 4**).

**Figure 4.**
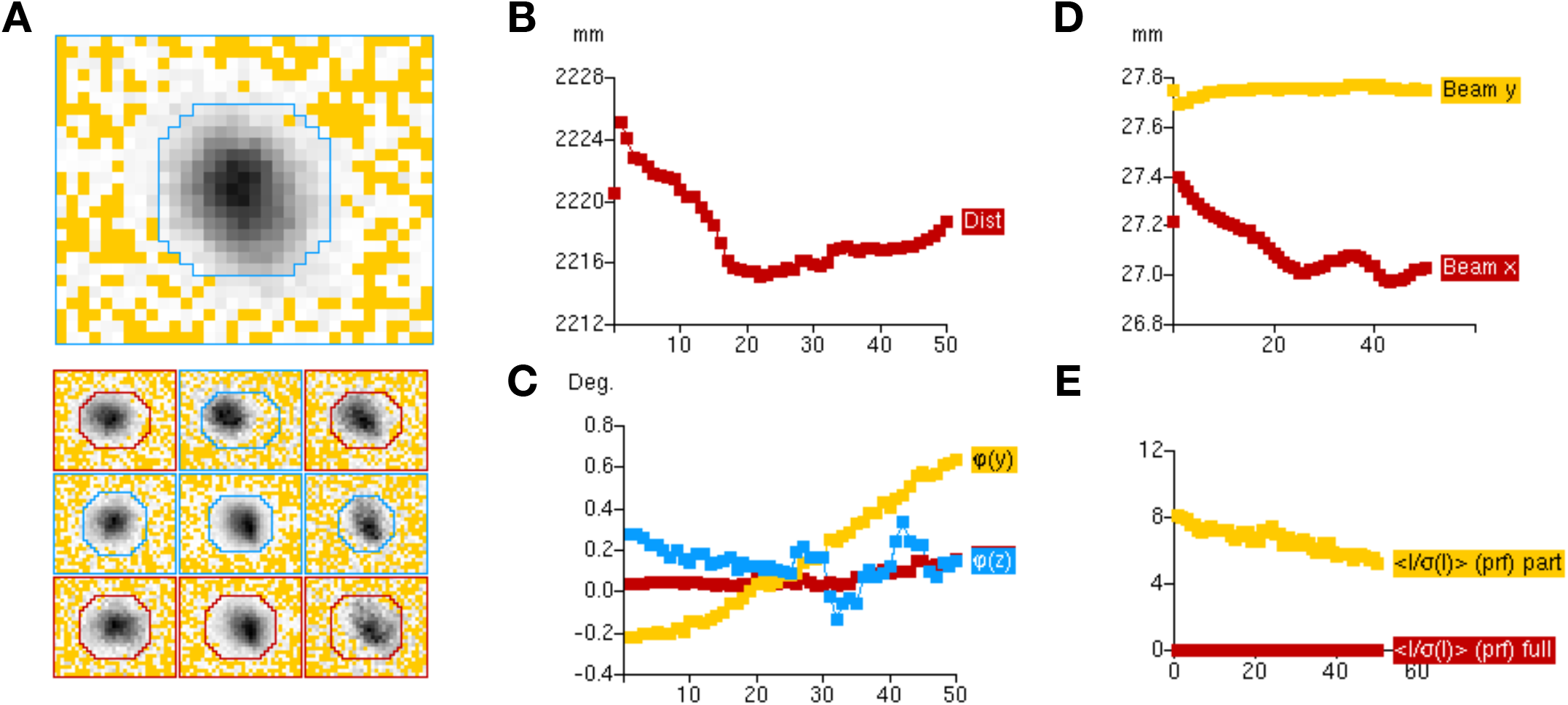
Statistic parameters during electron diffraction dataset processing by iMOSFLM. (A) A representative average spot profile of one diffraction image (up) and a representative standard profile for different regions of the detector (down). The red line indicates the profile is poor and averaged by including reflections from inner regions. (B) The crystal-to-detector distance changes with different diffraction images. (C) The crystal orientation changes with different diffraction images. (D) The electron beam position changes with different diffraction images. (E) The averaged SNR of diffraction spots changes with different diffraction images. The yellow curve represents all partial spots and the red one for all full spots.

**Figure 5.**
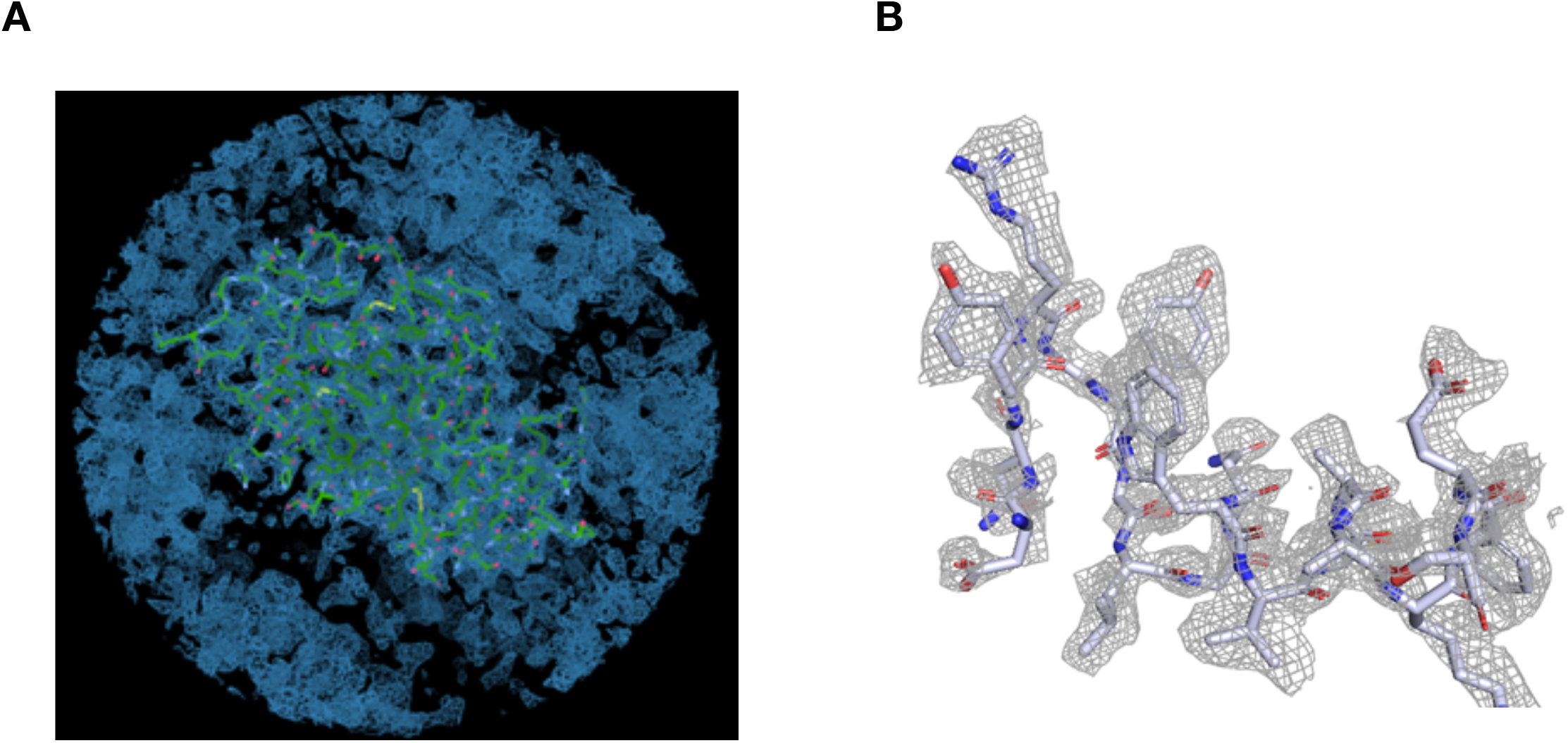
Electron potential map determined by molecular replacement. (A) The overall map fitted with the whole lysozyme structural model. (B) A zoomed-in view of the electron potential map around a selected region showing the resolution and quality of the map.

The profiles of diffraction spots suggested that the diffraction spots can be properly indexed, predicted and integrated (**Fig. 4A**). We found a significant jump of crystal-to-detector distance at the first few frames (**Fig. 4B**), which might be due to the precalibration error of the camera length. While, the detected variations of the crystal-to-detector distance and the crystal orientation (**Figs. 4B and 4C**) suggests that the stability of the microscope mechanical stage needs to be further optimized for MicroED experiments. From the statistics of the data processing (**Fig. 4D**), we also observed significant shift of electron beam position, which needs to be further investigated to find the reason. The existence of electron radiation damage and decay of crystal lattice could be indicated from the reduced SNR of diffraction spots during data collection (**Fig. 4E**). Finally, we collected seven datasets and merged into one dataset with space group of P4_3_2_1_2, the resolution of 2.5 Å, and the overall cumulative completeness of 94.0% (**Table 1**). The relative high R_merge_ of 0.356 would be caused by the inaccuracy of reciprocal spot intensity measurement, which could be most probably due to the poor hardware of our camera.

The final merged and scaled intensity data can be directly used for molecular replacement with a single significant solution. The calculated electron potential map based on electron scattering factor and the refined structural model shows a clear envelope of the molecule in the crystal (**Fig. 4A**). The resolution and quality of the map can further be reflected by the unambiguity of residue assignment in **Fig. 4B**. The final structure was refined to 2.5 Å with R_work_ of 35.9% and R_free_ of 40.0% (**Table 1**). Again, we believe that the relative high R value is due to the inaccuracy of reciprocal spot intensity measurement. Considering the current resolution of 2.5 Å, we did not intend to assign water molecule in the map.

Overall, in the present study, we developed a new approach to efficiently prepare suitable crystals for MicroED experiment. The *in situ* on-grid crystallization method avoids the potential loss of crystals during sample transfer, the single-side blotting method prevents the potential damage to the fragile crystals during vitrification, and the cryo-FIB fabrication method allows the crystals with the size of a few microns to be thinned for MicroED experiment. Finally, the large area of crystal cryo-lamella would yield a strong electron diffraction with good SNR. Thus a few crystals are enough to merge into a complete dataset for structural determination. Our work will greatly expand the availability space of crystals suitable for MicroED and fill up the gap between MicroED and X-ray crystallography.

In the future, we will further systematically investigate the influence of the thickness of crystal cryo-lamella for the MicroED data quality, and study whether the current and energy of focused ion beam would induce observable damage of crystal, which eventually affects the diffraction ability of the crystal.

## ACKNOWLEDGMENTS

We would like to thank Ping Shan and Ruigang Sui (F.S. lab) for their assistances. This work was supported by grants from Chinese Academy of Sciences (ZDKYYQ20170002 and XDB08030202) and the Ministry of Science and Technology of China (2017YFA0504700 and 2014CB910700). All the EM works were performed at Center for Biological Imaging (CBI, http://cbi.ibp.ac.cn), Institute of Biophysics, Chinese Academy of Sciences.

## Compliance with Ethical Standards

All authors declare that they have no conflict of interest. All institutional and national guidelines for the care and use of laboratory animals were followed.

## Human and animal rights and informed consent

This article does not contain any studies with human or animal subjects performed by any of the authors.

## Open Access

This article is distributed under the terms of the Creative Commons Attribution 4.0 International License (http://creativecommons.org/licenses/by/4.0/), which permits unrestricted use, distribution, and reproduction in any medium, provided you give appropriate credit to the original author(s) and the source, provide a link to the Creative Commons license, and indicate if changes were made. !

---

Movie S1. A representative movie showing the electron diffraction pattern changes along the rotation of the crystal.

## Supplementary Information

A SerialEM script to perform automatic collection of electron diffraction images from a continuously tilted crystal. (X denotes the starting angle, and Y denotes the end angle. Delay 1 means the interval time between each diffraction. 0.2 indicates that each rotation increases by 0.2°.)

~~~
=========================
start_angle = X
end_angle = Y
angle = $start_angle
Loop 1000
TiltTo $angle
Delay 1
ReportTiltAngle
R
S
angle = $angle + 0.2
If $angle > $end_angle
   break
Endif
EndLoop
=========================
~~~

